# Virtual reality for animal navigation with camera-based optical flow tracking

**DOI:** 10.1101/579813

**Authors:** Ivan Vishniakou, Paul G. Plöger, Johannes D. Seelig

## Abstract

**Background:** Virtual reality combined with spherical treadmills is used across species for studying neural circuits underlying navigation.

**New Method:** We developed an optical flow-based method for tracking treadmil ball motion in real-time using a single high-resolution camera.

**Results:** Tracking accuracy and timing were determined using calibration data. Ball tracking was performed at 500 Hz and integrated with an open source game engine for virtual reality projection. The projection was updated at 120 Hz with a latency with respect to ball motion of 30 ± 8 ms.

**Comparison with Existing Method(s):** Optical flow based tracking of treadmill motion is typically achieved using optical mice. The camera-based optical flow tracking system developed here is based on off-the-shelf components and offers control over the image acquisition and processing parameters. This results in flexibility with respect to tracking conditions – such as ball surface texture, lighting conditions, or ball size – as well as camera alignment and calibration.

**Conclusions:** A fast system for rotational ball motion tracking suitable for virtual reality animal behavior across different scales was developed and characterized.

## 1. Introduction

Virtual reality (VR) is used across species for studying neural circuits underlying behavior [1]. In many implementations, animals navigate through virtual realities on a spherical treadmill – a ball which can be freely rotated around its center of mass [1, 2, 3, 4, 5].

Tracking of ball rotation is typically accomplished using optical mice which are based on low-resolution, high-speed cameras integrated with a light source for measuring displacements when moving across a surface. Movement across the surface results in optical flow – the displacement of features across the camera sensor. Such features can for example be speckle-like reflections from surface roughness; comparing these speckle images between different frames then allows computing the displacement using hardware-integrated image processing.

Optical mice come however with some limitations for measuring ball rotation. First, a single optical mouse measures displacements only in two directions and therefore two mice are required for tracking all three degrees of freedom of ball motion. Secondly, limited or no control over the onboard processing algorithms as well as camera settings requires careful calibration. In particular, if the mouse sensors can’t be placed in direct proximity of the ball surface, accurate alignment of the two sensors as well as calibration with respect to surface properties and lighting conditions is necessary [5]. As an approach that overcomes some of these limitations, real-time tracking was developed with a single high-resolution camera for situations where a uniquely patterned ball can be used [6]. In that case, ball orientation was calculated by matching each recorded frame to a map of the entire ball surface pattern. Using a high-resolution cameras allows control over all recording parameters and offers the freedom to choose a custom algorithms and test its performance with simulated data [6].

Here, we combine the flexibility of optical flow-based tracking with the advantages of using a high-resolution camera. The developed system tracks optical flow of rotational ball movement at 500 Hz using a single camera and is integrated with an open source virtual reality environment and two projectors [7] with a refresh rate of 120 Hz [8] and overall latency of 30 ± 8 ms – from detecting ball movement to projecting the updated virtual reality image. This is achieved using off-the-shelf hardware components and open source software. The system is easy to align and calibrate, can be used under different conditions and at different scales, and can be integrated with two-photon imaging or electrophysiology.

## 2. Materials and Methods

### 2.1. Virtual reality setup components

A schematic of the setup is shown in Figure 1. Two digital micromirror devices (DMD, DLP LightCrafter 6500 by Texas Instruments) were used for projecting the VR onto an angled screen from two sides [8, 7]. The projectors are FullHD digital micromirror devices with HDMI and DisplayPort interfaces allowing them to display images the same way as standard monitors. In our setup, the DMD was set to use DisplayPort input and display the frames at a maximum frame rate of 120 Hz (see Appendix for details). This frame rate can only be achieved when frames are displayed with bit depth 1 (binary images). To display different gray levels, a dithering technique is used and the density of white pixels is varied proportionally to the brightness level of the displayed region with an ordered Bayer matrix [9] (see Appendix for details). Two collimated blue LEDs (M470L3, Thorlabs) were used as light sources. Light reflected off the DMD was projected from the outside (seen from the fly’s perspective) onto a screen made from black paper.

**Figure 1:**
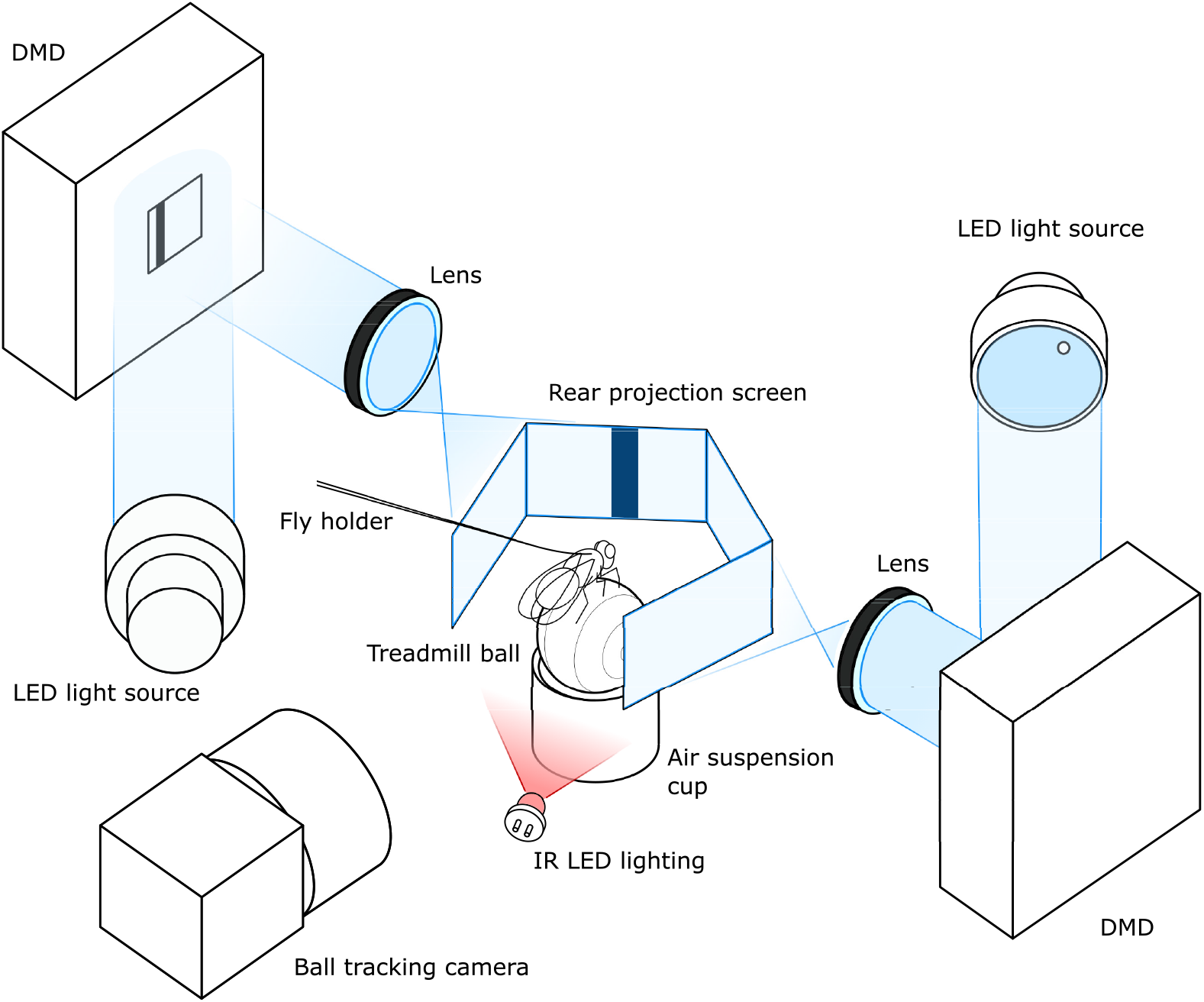
Schematic of behavior setup for *Drosophila melanogaster.* A 6-mm polyurethane ball is air-suspended in a holder and serves as an omnidirectional treadmill. The fly is glued to a thin metal wire. A rear projection screen is used for displaying the virtual reality (VR) in the fly’s field of view. Two DMDs, LED light sources, and two projection lenses are combined to project from two sides onto the screen. A single tracking camera is set up to capture ball images at the rate of 500 fps. Infrared LED lighting invisible to the fly is used for illuminating the ball.

The spherical treadmill used for the current experiments is described in [5]. Briefly, a 6 mm diameter polyurethane foam ball is held in a cup-shaped holder and suspended in an air stream. The ball is illuminated with two IR LEDs and monitored with a camera. A *Basler acA6Ą0-750um* camera was used for tracking. The camera uses a USB 3 interface to communicate with the PC and the bandwidth of 350 MB/s allows transfer of full-frame images (640 × 480 × 8bpp mono color) at a rate of 751 frames per second (fps). For tracking, a frame rate of 500 fps was used, with an exposure time of 100 microseconds under infrared illumination. The resolution was 224 × 140 pixels which results from cropping and 2 × 2 binning of the full frame. Binning (summing the signal from adjacent pixels into a single one) increased signal-to-noise ratio and reduced the image readout and transfer time. A Computar camera objective M2514-MP2 F1.4, f = 25mm together with a Computar EX2C extender were used for imaging the ball [5].

### 2.2. Integration with virtual reality

For virtual reality applications a 3D rendering game engine (Urho3D, an open-source C++ game engine [10]) was used and the x, y- and z-rotations of the ball were mapped to planar motion (forward-, sidestepping- and turning components) of the animal in the virtual reality (this was done similarly to [6], see Appendix for details).

The application is designed to run ball tracking and environment rendering in two separate execution threads. The image processing thread is a loop acquiring camera frames and calculating ball displacements, tuned to run one iteration in under 2 ms, resulting in a tracking frequency of 500 fps. The rendering thread runs the game engine, which models a virtual environment in 3D.

Setting up the virtual environment requires specifying the arena geometry as well as lighting and rendering properties. Further, the virtual cameras have to be configured such that the virtual environment is rendered form the angles corresponding to the position of the screen. Additional parameters are required to describe the mapping of ball rotations to movement in the virtual environment; these transformations take into account the size of the ball and the orientation of the tracking camera relative to the forward direction of the animal. This is all done with the in-built Urho3D engine’s scripting language which also describes each environment in a separate script file, containing instructions about which objects are to be created in the scene. In each update of the game engine the ball tacking signal is interpreted as the displacement of the animal, and the virtual camera positions and orientations are updated before being rendered and displayed on the screen.

Other game engine capabilities used in the application are dithering shader to output binary frames suitable for the DMD and networking to broadcast the state of the VR to any client, for example for monitoring the VR or a script that triggers any other external stimuli. The application can also be used without visual output for tracking the animal’s activity. The configuration of the virtual environment is described in detail in the Appendix.

The tracking data is saved in a text log file: for each camera frame a new line is added with the timestamp of the frame, x-, y-, z-ball displacements and x-, y-, z-position of the animal in the virtual environment. Additionally, the tracking camera is set to output a frame trigger pulse with each recorded frame, which can be used for synchronization with other applications, such as two-photon imaging.

All experiments were performed on a PC with an Intel Xeon E5-1620 v3 @ 3.50GHz (8 cores) CPU, 32Gb DDR 4 @ 2993MHz of RAM, a NVIDIA Quadro K620, 2Gb DDR3 RAM, 384 CUDA cores GPU and Microsoft Windows 8.1 Enterprise edition operating system. OpenCV 3.4.1 and opencv-contrib module were compiled using Microsoft Visual Studio Toolkit 14.0 (VS2015) in release configuration with CUDA 9.1 for tests of the GPU-accelerated algorithms.

## 3. Theory/calculation

### 3.1. Modeling of optical flow camera projections

In this section a model is developed that describes how optical flow generated by ball rotations around different axes is projected onto the camera sensor. Predictions of this model will be used as fit functions to extract ball rotation parameters from measured optical flow distributions.

Ball motion is described by specifying an axis of rotation and an angular velocity (axis-angle representation [11]). Assuming a Cartesian coordinate system *O* with the origin at the center of the ball and a corresponding ISO-conventional [12] spherical coordinate system S as shown in Figure 2, a point *p* on the surface of the ball can be described with a vector **p**_*S*_ = (*r_ball_, θ_p_, φ_p_*). If *O* is selected so that its z-axis coincides with the axis of ball rotation, it becomes a convenient parametrization since only the azimuthal angle is varying with time if the ball rotates around the polar axis with angular velocity *ω*:

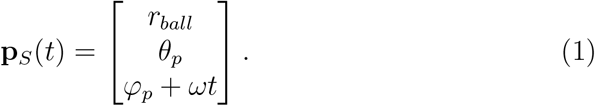

**Figure 2:**
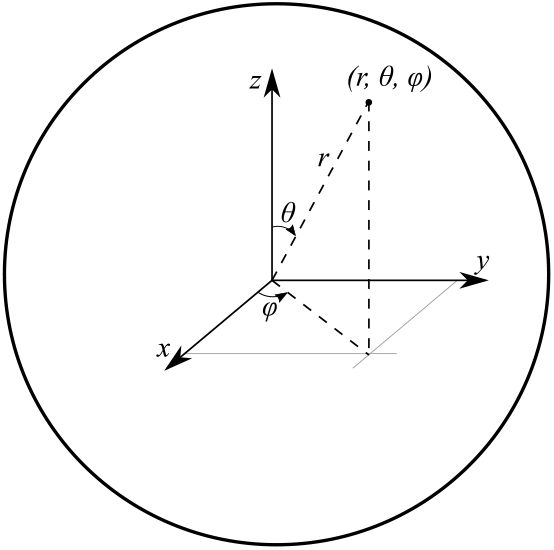
Spherical coordinate system complying with the ISO 31-11 convention: a point is addressed by radius *r*, polar angle *θ* and azimuthal angle *φ*

In the Cartesian coordinate system *O* the same trajectory is expressed as

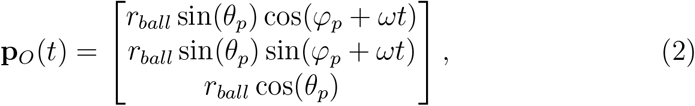

where *θ_p_* and *φ_p_* are the polar and azimuth angle of a point on the surface of the ball and *r_ball_* is the constant radius.

Using a pinhole camera model [13] we can express the projection of points on the ball surface onto the camera sensor. We here follow the same naming and frame orientation convention as the OpenCV library [14, 15], shown in Figure 3. The trajectory of a point on the ball surface in the camera frame *C* is

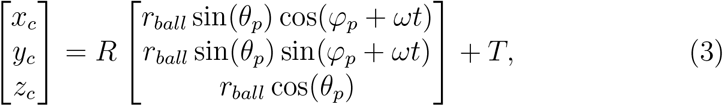

where *R* is the rotation matrix representing the orientation of the ball’s axis of rotation and *T* is the position of the ball’s reference frame origin in the camera frame *C*. The image plane coordinates (*u, v*) of the projected point *p*′ are expressed as

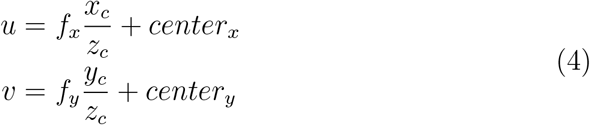

where *f_x_* and *f_y_* are the focal lengths expressed in pixel units and (*center_x_, center_y_*) is the image center. The ball is centered in the camera’s field of view and therefore *T* = [0, 0,*d*], where *d* is the distance between the center of the ball and the camera aperture.

**Figure 3:**
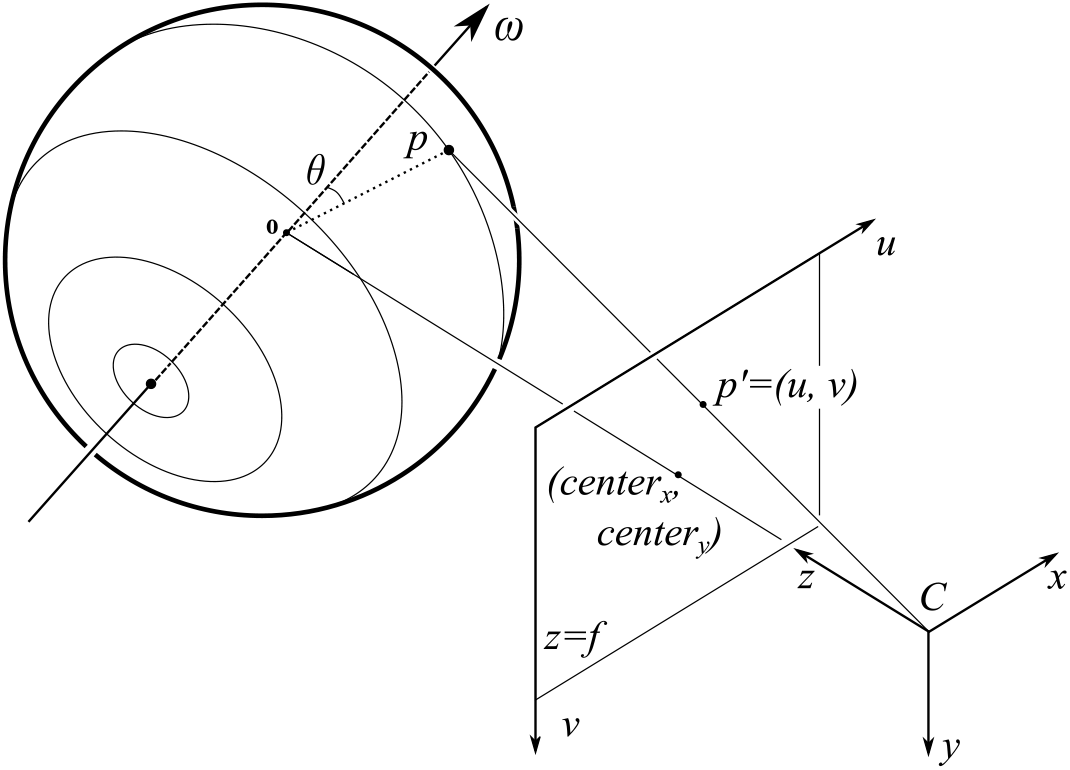
Projection of points on the ball with a pinhole camera model. Point *p* on the ball surface can be described with coordinates (*r, θ,φ*) in a polar coordinate system aligned with the ball’s instant axis of rotation. According to expression (3) it can be expressed in Cartesian coordinates (*x, y, z*) in the cameras reference frame C and projected onto the point *p*’ = (*u, v*) in the image plane according to equation (4).

By calculating the ball point projection between two consecutive frames at time t and t + Δt, with Δt equal to the frame period, one can calculate the displacement of the point projection and predict optical flow. If all camera parameters as well as ball location and ball size are known, the model can be used to calculate the optical flow distribution on the camera. The model was implemented using the symbolic math library Sympy.

The angular velocity vector can be expressed as a sum of three orthogonal angular velocity components. For a camera-centered ball, it is convenient to choose the axes of a frame aligned with the camera frame with its origin at the ball’s center. This gives three distinct rotations which can be registered by the camera as x-, y- and z-rotations (see Figure 3).

Filling in the matrix R in expression (3) to obtain ball rotations around the x, y, and z camera frame axes, respectively, and combining it with (4) yields the trajectories of the ball’s point projections on the camera (for rotation around the specified axis):

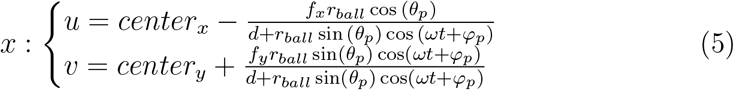

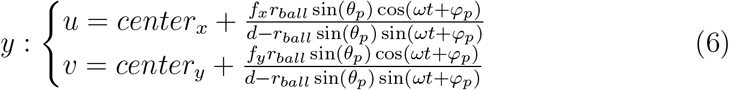

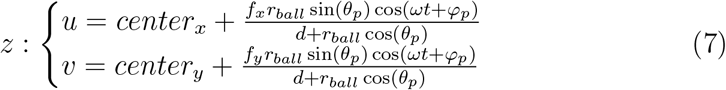

As seen in Figure 4 a-c, rotations around the three different camera frame axes produce recognizably different velocity field distributions. To distinguish them, a ring-shaped region of interest is selected around the center of the ball (Figure 4, d-f). For each point in the ROI, two principal directions are selected (radial – orthogonal to the ROI circle at this point – and tangential to the ROI circle at this point, directed counterclockwise). The optical flow vector is then projected onto these directions to find its radial and tangential components with respect to the ROI (Figure 4, g-i). These radial and tangential components of the optical flow build distinct distributions over the ROI, which allows to separate the motion components (Figure 4, j-l).

**Figure 4:**
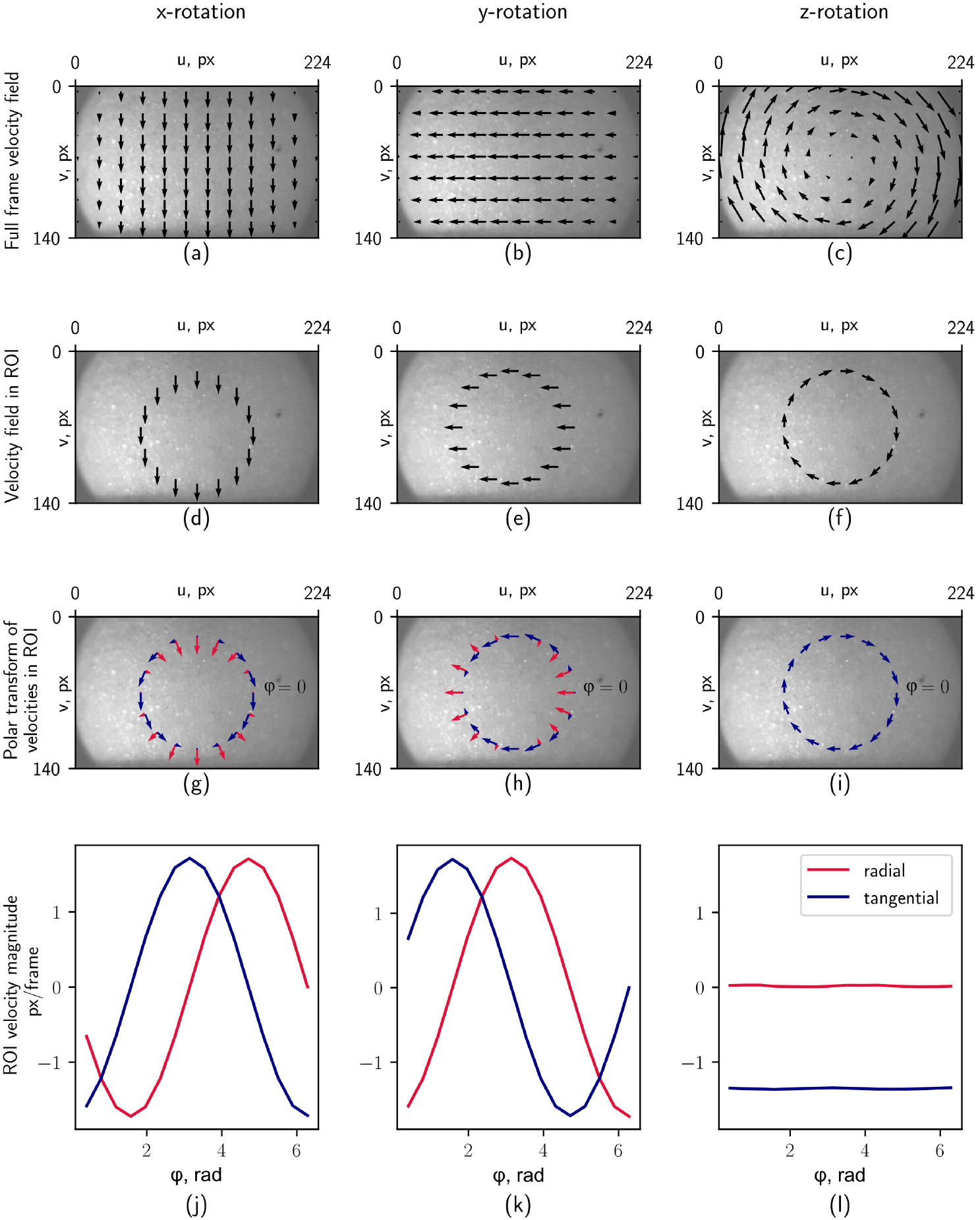
Predicted optical flow produced by rotations of the ball around the respective three independent axes of rotation.

These distributions can be fitted well by the following two functions

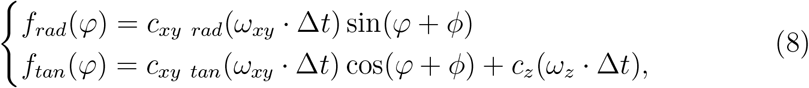

where *c_xy rad_* and *c_xy tan_* (px/rad) are calibration factors relating the angular displacement for angular velocity vectors lying in the xy-plane to the respective optical flow (see Appendix for details). Similarly, *c_z_* (px/rad) is a factor relating angular displacement for angular velocity vectors lying on the z-axis to optical flow; (*ω_xy_* · Δ*t*) and (*ω_z_* · Δ*t*) are the angular displacements about a rotational axis lying in the xy-plane or on the z-axis, respectively, and *ϕ* is the orientation of the axis of rotation in the xy-plane. Note, that radial and tangential components of optical flow induced by *ω_xy_* have different coefficients, since the magnitudes of the respective optical flow distributions do not match, while the *ω_z_* rotation induces only tangential optical flow, and a single coefficient *c_z_* is sufficient. The calibration factors subsume all setup parameters, such as camera focal length, and can be determined as described below and in the Appendix.

### 3.2. Finding calibration factors by function fitting

Calibration factors were determined experimentally by recording a sequence of frames with two cameras pointed at the ball center and positioned orthogonally to each other (see Figure 14 in the Appendix). The z-component is a planar (that is, in the camera focal plane) motion which can be measured accurately with optical flow (as was verified using simulated as well as motor actuated ground truth data), the xy-coefficient can be found by regression between the estimates of the ball motion based on the two cameras. This is described in more detail in the Appendix.

### 3.3. Tracking algorithm

The ball tracking algorithm takes two consecutive frames of ball motion and calculates the three independent rotational displacements of the ball, i.e. the x-, y- and z-axis angular displacements, by fitting their projections onto the radial and tangential directions with function (8) in a circular ROI. Since the distribution of optical flow in the circular ROI is very noisy the optical flow is averaged over a band (limited by a minimal inner and maximal outer ring, see Appendix for how ROI size was determined) in the radial direction to increase robustness. This results in the following algorithm:

**Data**: Two grayscale images of the ball video

**Result**: Angular velocity vector (*ω_x_, ω_y_, ω_z_*)

**Input**: *frame*_1_, *frame*_2_

1. polar transform *frame*_1_, *frame*_2_ with respect to the center of the ROI
2. crop the polar-transformed *frame*_1_, *frame*_2_ to keep only the ROI
3. *flow_u_, flow_v_* ← compute_optical_flow(*frame*_1_ (*cropped*), *frame*_2_ (*cropped*))
4. *flowROI_rad_, flowROI_tan_* ← average *flow_u_, flow_v_* over radius
5. fit *flowROI_rad_, flowROI_tan_* with function (8) by finding (*ω_xy_* · Δ*t*), *φ*, (*ω_z_* · Δ*t*)
6. (*ω_x_, ω_y_, ω_z_*) ← *ω_xy_* sin(*ϕ*), *ω_xy_* cos(*ϕ*), *ω_z_* **return** (*ω_x_, ω_y_, ω_z_*)

Data from the algorithm’s intermediate steps are shown in Figure 5. The polar transform (Figure 5, b) is calculated before computing optical flow, allowing the following optimizations: first, the ring-shaped ROI transforms into a rectangular image section, where one dimension corresponds to the azimuthal angle of the ROI *φ* and the other to the radius *ρ*. This means that the ROI can be isolated by cropping the image and optical flow can be calculated for a small fraction of the full frame. Secondly, in the polar-transformed ROI, the optical flow *u* and *v* components correspond to the radial and tangential motions directly, which eliminates the need of flow re-projection. The dependence of the optical flow calculation on the ROI position which results from the polar transform (see Figure 5, b) is included in the experimentally determined calibration coefficients.

**Figure 5:**
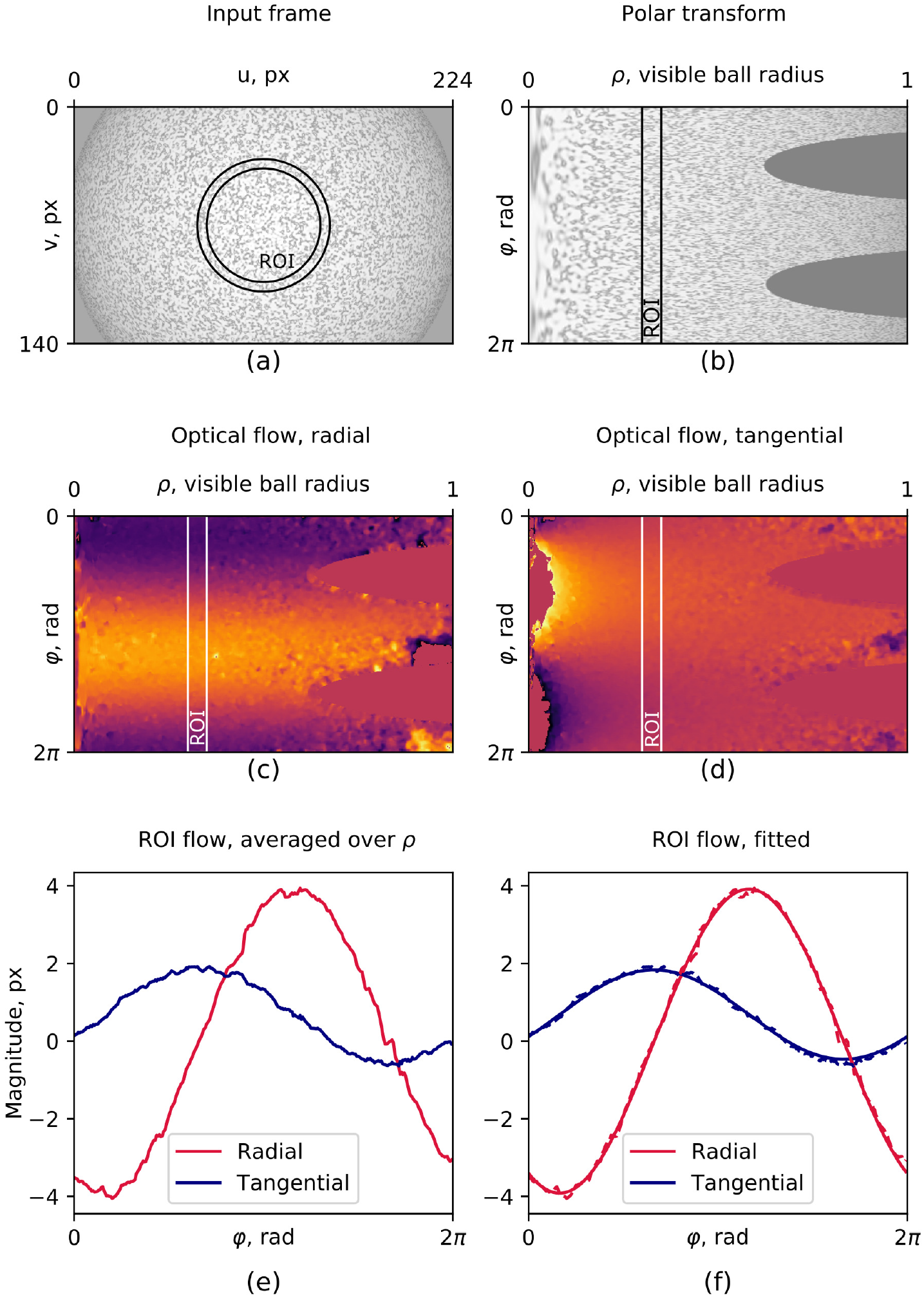
Ball rotation evaluation from optical flow. Input frame (a) is polar-transformed (b) and optical flow (c, d) is extracted by comparing it to the previous frame. The distribution of optical flow in the ROI (e) is then fitted with function (8) (f).

Other optimizations included passing over the polar-transformed and cropped frame to the next iteration to save on those operations, and using the previous iteration’s function fitting result as a prior for the solver in the next frame. Function fitting is done with Downhill Solver included in OpenCV, which implements the Nelder-Mead downhill simplex method [16].

For a compiled executable of the VR application please send an email to.

## 4. Results

### 4.1. Tracking algorithm implementation

For real-time applications the optical flow calculation was implemented in a C++ program using OpenCV 3.4.1. Best performance in terms of processing speed and tracking accuracy was achieved with the Farneback optical flow algorithm on a 60 × 140 ROI scaled down to 30 × 140 (see the Appendix for a comparison of a range of optical flow algorithms and for selecting ROI parameters). Preprocessing (polar transform, cropping and resizing) together with optical flow estimation took 1.45 ms (standard deviation s = 0.54 ms), the function fitting by downhill solver was capped at 30 iterations to keep its run-time under 0.4 ms. These parameters allow to achieve real-time tracking at 500 frames per second with the current PC configuration. All time measurements were performed with the cv::TickMeter class.

We noticed that occasionally frames were dropped, for example when starting the program or due to other operating system related processes. Such frame drops can be detected in the saved data file by looking for time stamps with a separation of more than 2 ms. If a frame is dropped, the next displacement is calculated with the latest available frame before the frame drop.

### 4.2. Evaluation of tracking accuracy

The evaluation of tracking accuracy is based on comparing ground truth *ω*_GT_ and estimated angular velocity *ω*_est_. Three metrics are selected for this comparison: the absolute error (in degrees per frame)

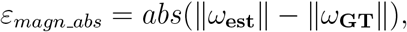

the relative (percentage) magnitude error

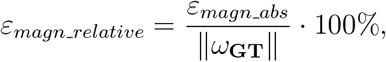

and the orientation error (in degrees), which is the angle between the ground truth and estimated rotation vectors in the plane spanned by these two vectors,

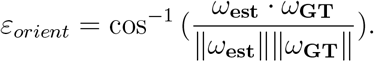

Two ground truth datasets were generated for testing the tracking performance (see Appendix for details). One consisted of simulated data which offered control of all rotation parameters, while however not fully replicating surface texture or lighting and camera conditions encountered in the actual experiment. A second dataset was generated by rotating a 60 mm ball (10x scale model for the 6 mm ball used for fly behavior) with a stepper motor.

Using the simulated data with accelerated rotations the operation range of the tracking alg
orithm was determined (see Appendix for details). Figure 7 shows the comparison of the integrated tracked trajectory with ground truth data. A value of 50^°^ error for the x-axis corresponds to a 2.6 mm error in a planar trajectory. The tracking error depends on the rotation speed of the ball (for the ROI and parameters used here, see Figure 15 in the Appendix). The average tracking error stays under 10% (magnitude) and 7.5^°^ (orientation) for rotations up to 1.70^°^ per frame, which corresponds to 45 mm/s linear speed for a 6 mm diameter ball.

Using a stepper-motor a ball was rotated at constant speed around a single axis (see Appendix for details). The integrated rotation magnitude error and orientation errors for one rotation sequence are shown in Figure 6.

**Figure 6:**
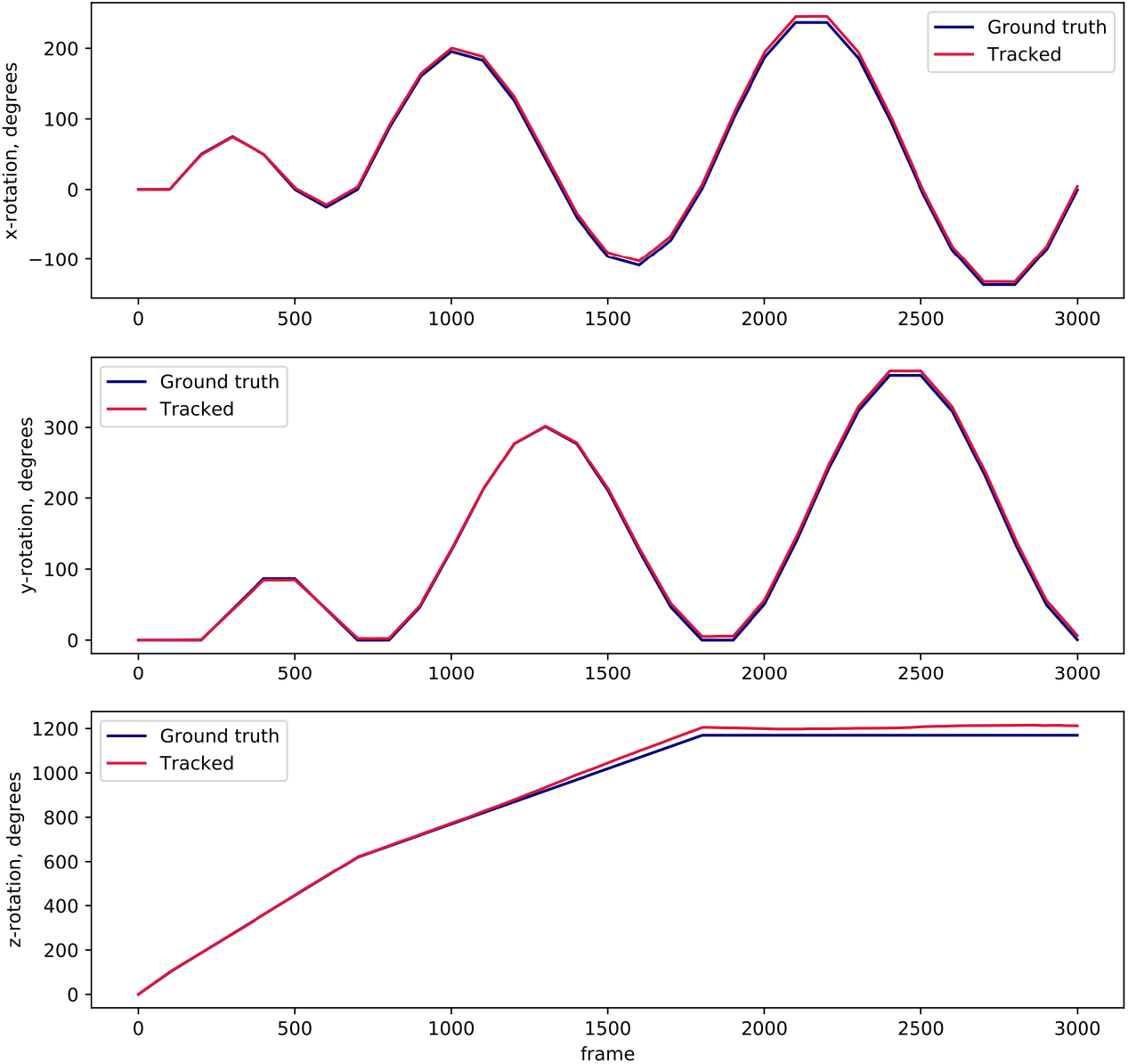
Integrated tracked trajectories and ground truth for stepping motor actuated ball movement at constant speed.

**Figure 7:**
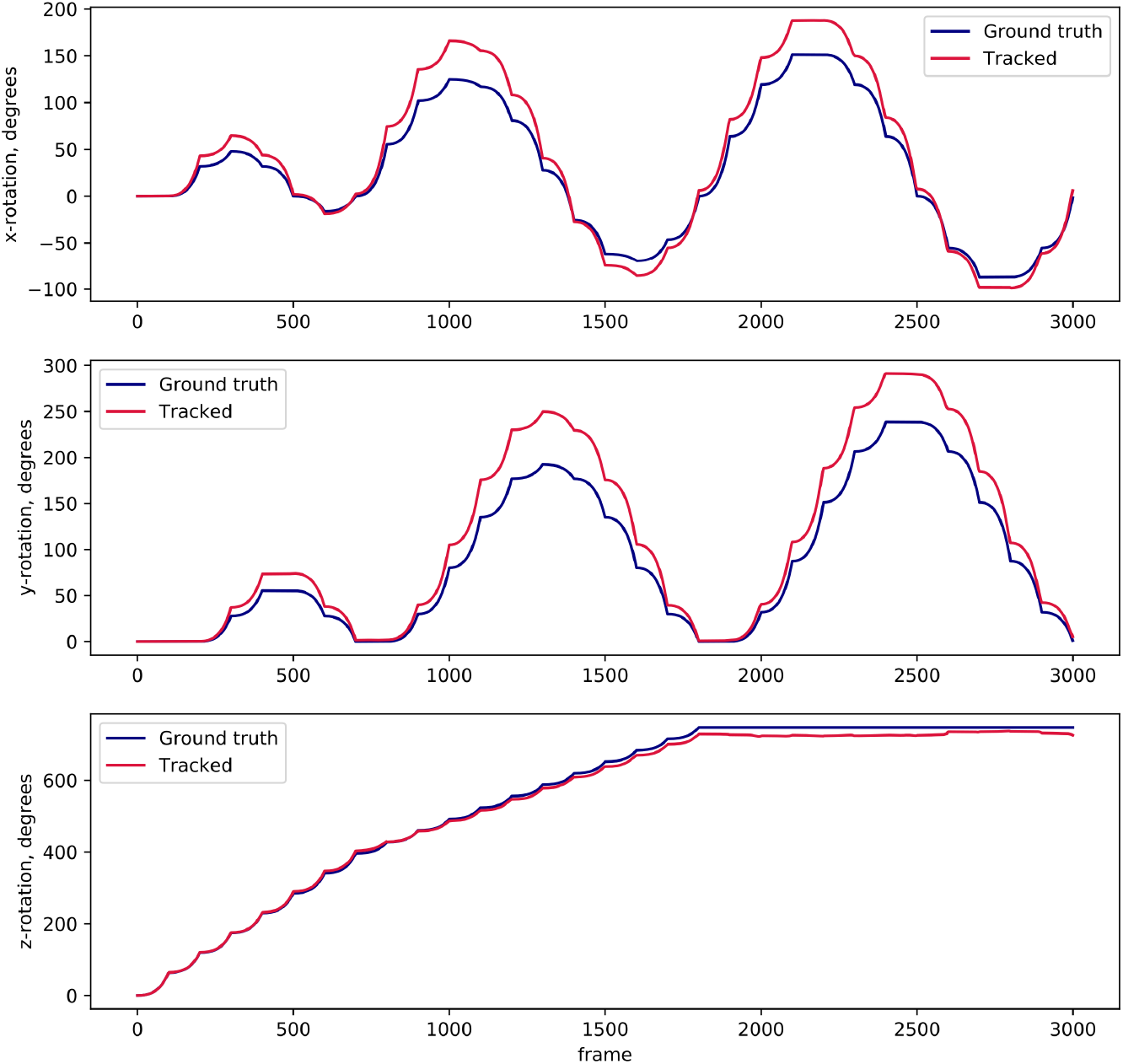
Integrated tracked trajectories and ground truth of ball motion for an example of simulated, accelerated rotations.

### 4.3. VR system timing

The run-time of the tracking algorithm was measured using internal timing. The total latency of the combined tracking and VR setup was estimated using a high-speed camera filming the VR display: the display was triggered to change state, and the state change was detected with the tracking camera. This sequence contains all the latency components for one cycle of a closed loop VR experiment.

The main component of the latency is the display device input lag. It depends strongly on the selected video mode (Table 1) and the lowest values were found at 120 Hz update rate with V-sync disabled.

**Table 1:** Latency measurement of a camera – virtual reality – display system

| Display mode | Median latency, ms |
| --- | --- |
| 120 Hz | 25 ± 8 ms |
| 120 Hz, V-sync | 33 ± 8 ms |
| 60 Hz | 50 ± 16 ms |
| 60 Hz, V-sync | 50 ± 16 ms |

### 4.4. Behavior experiments in VR

For testing the setup for fly behavior, we used a simple VR with a single bright stripe of 10 degree width and 45 degree height projected onto a two-part screen similar to [7] of 9 mm height and 2 times 18 mm width made out of black paper. The fly’s head was fixed to its thorax with UV glue, and the wings were glued to the tethering pin using UV glue to encourage walking behavior [17]. The stripe was visible for 4/5th of the flies virtual surroundings and disappeared for 1/5th behind the fly. This resulted in robust orientation behavior with the fly walking away from the stripe (see Figure 8).

**Figure 8:**
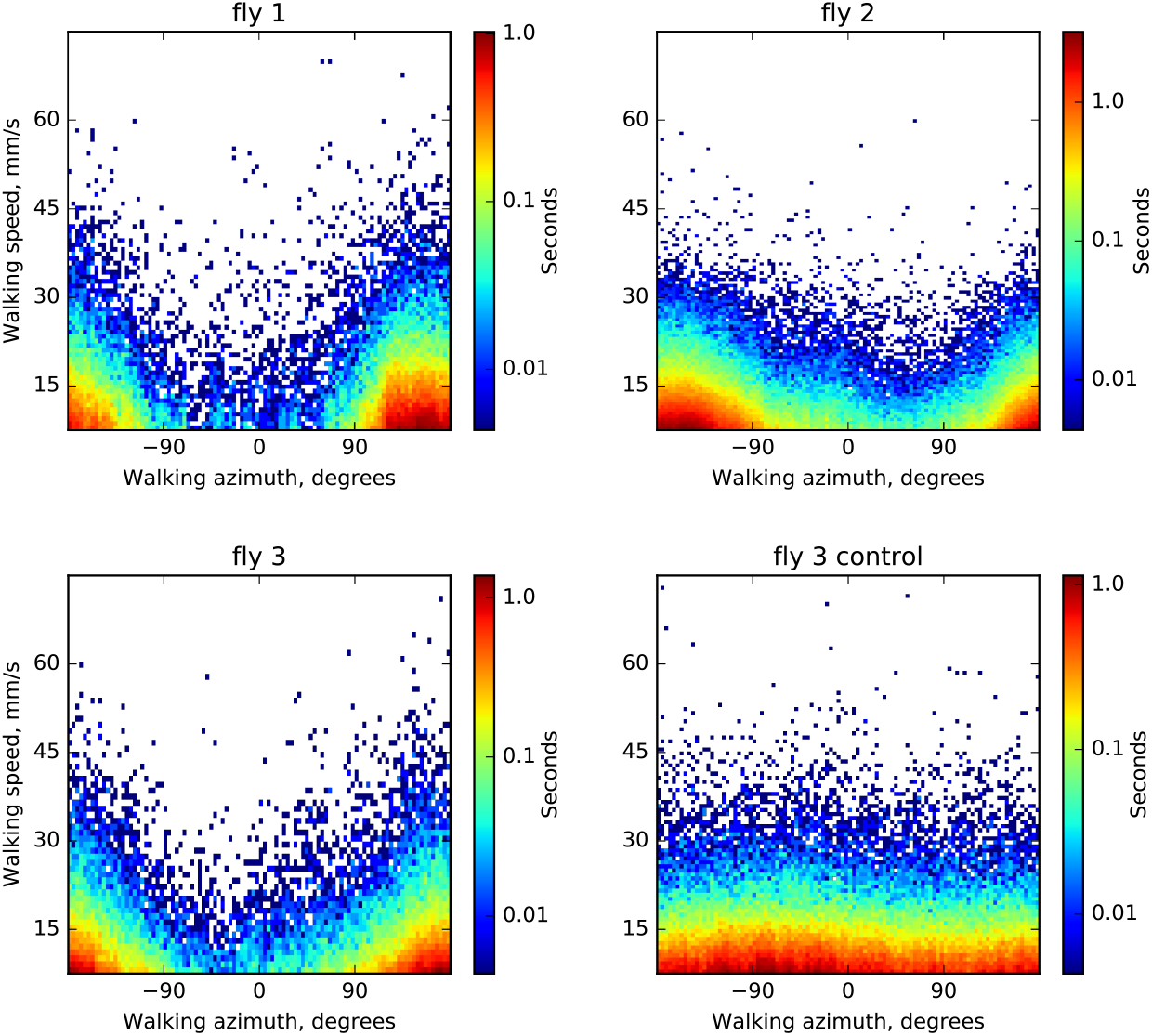
Walking speed and walking direction of three flies walking in closed loop with a bright stripe on a black screen (see text for details). Color indicates histogram bins (binned with a grid of 100 × 100 pixels) of time spent in each state for a total trial length between 15 and 25 minutes (depending on the fly)). In darkness the flies don’t show a pronounced orientation preference (as shown for fly 3, control).

## 5. Discussion

An optical flow-based method for tracking of spherical treadmill motion using a single high-resolution camera was developed and integrated with virtual reality projection. Treadmill motion was monitored at 500 Hz, the display update rate was 120 Hz, and the total latency for an entire VR cycle was 30 ± 8 ms (from reading out ball movement to projecting the updated VR frame on the display). The system combines the flexibility of optical flow based tracking with the convenience of using a single high resolution camera, off-the-shelf components, and open source software.

Compared with an optical mouse based system the approach developed here offers independent control over the different components of the system, such as camera settings and optical flow processing algorithm. Image processing is therefore not limited to a preset and unknown algorithm, but a suitable approach can be selected and its parameters can be tuned (see Appendix for a comparison of all optical flow algorithms tested). Additionally, the resulting tracking accuracy and the operation range (which for all optical flow based methods depends on the combination of field of view, movement speeds and processing algorithm) can be tested using simulated data. Different from approaches using optical mice, tracking of all three degrees of freedom of ball rotation is achieved with a single camera which can be positioned flexibly and can easily aligned and calibrated. This also facilitates adapting the system to a variety of experimental conditions and scales.

As an alternative to optical flow approaches, a solution relying on a unique patterning of the ball surface was implemented in [6]. While this pattern matching approach avoids integration errors, optical flow can on the other hand track any surface with sufficient speckle contrast under a variety of lighting conditions and can be adapted to different ball sizes.

Compared to cameras, optical mouse sensors typically have a higher frame rate (at the cost of lower pixel resolution) and time jitter and latencies are shorter on a dedicated processing chip. For example, [3] measured the temporal accuracy of a customized optical mouse sensor and found a tracking latency of less than 500 *μs*, compared to 2 ms here. However, as the latency time breakdown shows (see Table 2), the delay is mostly caused by the standard display interface (DisplayPort). Since the camera tracking latencies and jitter are small compared to the display latencies and jitter, this should have a very limited impact on VR behavior. The overall latency of the closed-loop virtual reality system was 30 ± 8 ms on average, comparable to optical mice-based solutions and about three times faster than another camera-based pattern matching approach [6]. The display update rate of typical virtual reality setups with data transfer through the display port is generally limited to 60 to 120 Hz. Typical latencies found with such PC and camera-based solutions are between 50-90 ms [18, 6] and have been shown to be acceptable for closed-loop behavior experiments with a variety of species [1, 18]. We tested the system for behavioral performance in fruit flies and found a robust avoidance response of a bright stripe under the display conditions tested.

**Table 2:**
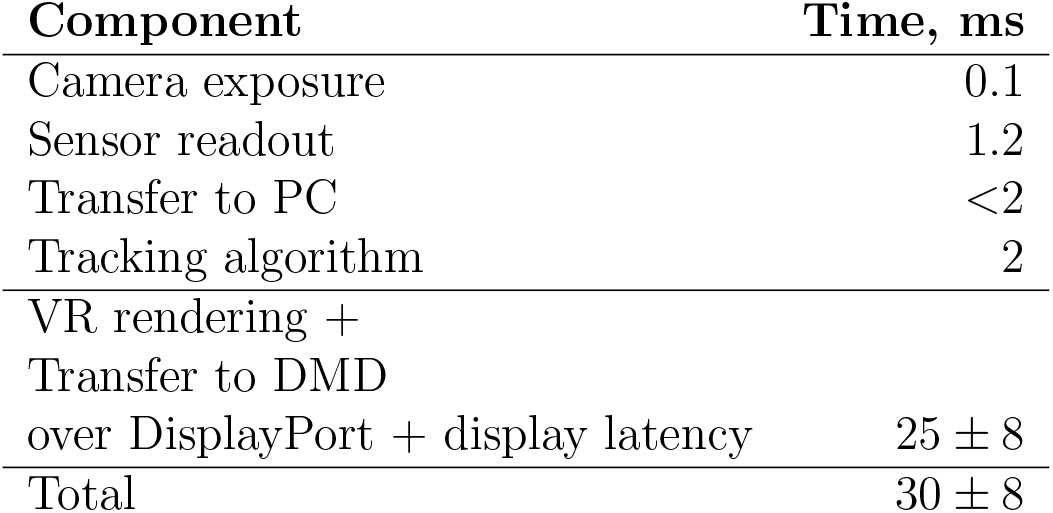
Latency time breakdown for the closed-loop virtual reality setup

Overall, a VR system based on optical flow measurements with a high-resolution camera was developed with short latencies suitable for VR experiments at different scales.

## 6. Acknowledgements

We would like to thank Santosh Thoduka for suggestions on camera calibration and helpful discussions, Fiona Ross and Stephanie Miceli for initial behavioral tests, Andres Flores for help with behavior analysis and designing hardware for stimulus projection, Bernd Scheiding for electronics support, and Dan Turner-Evans for help with IR illumination.

## 7. Appendix

### 7.1 Tracking settings

Tracking-specific settings are stored in the `Config\tracking.cfg` file and include the path to the camera configuration file, the name of the camera to be used for tracking, as well as calibration coefficients as explained in section 3.1. Those parameters can be found in the corresponding sections of the config file:

~~~
[camera]
LoadCameraSettings=true
CameraSettingsFile=Config\camera_settings. pfs
TrackingCameraName=ball_cam

[calibration]
BallCenterX = 112.0
BallCenterY = 70.0
BallRadius = 116.0
CxyRad = 100.31
CxyTan=76.85
Cz = 20.63
~~~

The camera was set to capture images at 500 fps, and the exposure time was set to 100 *μs* to prevent motion blurring (and required using sufficient infrared intensity). The resolution was set to 224 × 140, the highest resolution that was handled by our system in the available 2 ms per frame processing time. The settings of the camera can be saved in the Pylon configuration file and set to be loaded in the tracking configuration.

### 7.2. DMD settings

In order to use the LightCrafter 6500 DMDs at a 120 Hz display rate, they need to be connected through the DisplayPort interface and have to be configured to use Dual pixel clock mode, run in Video Pattern mode: this is an operation mode where each frame arriving through the video interface (RGB 8bpp) is regarded as 24 independent bit planes and each can be independently selected to be displayed with an arbitrary timing diagram. In this mode the most significant bit of the green channel is taken and displayed for 7 ms, resulting in 120 fps display rate. Additionally, it might be necessary to enable 120 Hz refresh rate in the Windows display adapter settings for both DMDs, which are visible to the operating system as normal displays.

The most convenient way to automate the DMD configuration is by saving these settings in a batch file (sequence of commands sent to the DMD) and upload it to the DMD as a startup script (this is a feature supported by LightCrafter 6500).

### 7.3. Virtual environment settings

Virtual reality has several configuration parameters which are specified in the `Config\vr.cfg`. It has game-engine specific parameters in the corresponding section, like the position of the VR window in screen coordinates and its size. Since we used two DMDs, it is spanned over twice the resolution of the DMD in width and located to the right of the main screen. `VSync` is disabled, since it is otherwise introduces additional latency to the display. `minFps` is set equal to the refresh rate of the displays; `maxFps` and `maxInactiveFps` are capping the update rate of the game engine. maxInactiveFps is the frame rate limit for a window while it is not in focus, i.e. when the user is working with other applications. This is almost always the case, since the game window is not visible for the computer user and other applications, for example for monitoring the VR, are used. The default value is 60 fps, and therefore needs to be increased in order to use the full capacity of the 120 Hz DMD displays.

The “transforms” section of the configuration defines how the ball rotation maps to motion in the virtual environment. The tracking signal is calibrated so that it presents angular displacements of the ball (in radians) about the x-, y-, and z-axes of the camera. Depending on how the camera is oriented relative to forward walking direction of the animal, the mapping varies. `ballXYZtoArenaXYZ` is a 3 × 3 rotation matrix, and

~~~
arena_displacement = ballXYZtoArenaXYZ · ball_displacement.
~~~

Similarly, the turning in the virtual environment is calculated from the ball displacement as a dot product with `ballXYZtoArenaYaw`:

~~~
arena_yaw_displacement = ballXYZtoArenaYaw · ball_displacement.
~~~

In the provided example the tracking camera is placed behind the ball and is aligned with the forward walking direction of the fly:

~~~
[transforms]
ballXYZtoArenaXYZ=0 0 – 3.0 0 0 0 3.0 0 0
ballXYZtoArenaYaw=0 – 57.3248 0
~~~

The 3 mm radius of the ball is incorporated in the motion transform, and the yaw rotation contains a conversion from radians to degrees.

### 7.4. Arena scripts

Although the Urho3D game engine allows saving and loading 3D environments in XML format, a more readable and convenient way is to declare environment properties in a script. Urho3D supports a compiled scripting language called AngelScript [19], which is object-oriented, and features syntax similar to C++. The scene is represented in the game engine as a hierarchical tree structure: each entity of the 3D environment is a node that can be attached to either other nodes or to the root node of the scene. The coordinates of the nodes are always interpreted in the coordinate frame of their parent nodes. In-detail information is provided in the game engine’s manual [20], and sample arenas are documented. Dithering, which is required for displaying grayscale images with a DMD operating in binary display mode is done with a postprocessing shader, which is added to the rendering pipeline of the game engine. The configuration of the VR display is also shown in a documented example script.

### 7.5. Ground truth datasets

One ground truth dataset was recorded with a 60 mm polystyrene foam ball (10 × scale model) mounted on a stepper motor [6]. The motor was controlled using Grbl firmware for Arduino boards [21]. The distance from the camera to the ball is was scaled accordingly 10 × as well. Sequences of 1000 frames were recorded with an angular step size of 0.281, 0.562, 1.124 and 1.686 degrees per frame, in two orientations corresponding to y- and z-rotations.

A second ground truth dataset consisted of simulated data which allowed full control over all parameters: a 3D scene was reconstructed with Blender 3D including a camera with focal parameters identical to the actual tracking setup, a light source and a ball with grainy surface texture. The model of the ball was rotated and rendered using Blender’s API and in-built Python interpreter. Two simulated rotation sequences were created: The first one was used for calibration and accuracy validation and consisted of rotations of 1^°^ per frame (corresponding to about half of the maximal walking speed of the fly), with 30 different rotation axes and 100 frames for each axis. The second dataset consisted of rotations of exponentially increasing magnitude from 0^°^ to 2^°^ per frame, intended to test the operating range of the tracking algorithm. The orientations of the rotational axes for both datasets were selected evenly distributed.

An actual tracking camera image is shown side-to-side with a simulated image in Figures 9 and 10: the real image has out-of-focus areas, some bright speckles from direct reflection of the lighting, and an overall uneven brightness distribution.

**Figure 9:**
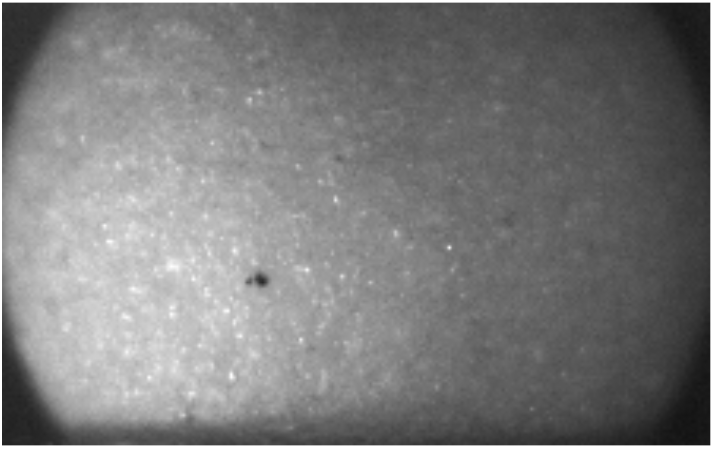
Frame captured by tracking camera in VR setup.

**Figure 10:**
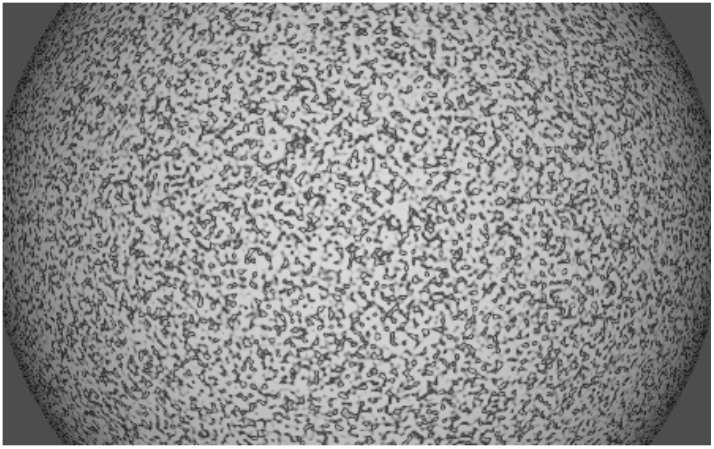
Frame from simulated ball rotation sequence.

### 7.6. Optical flow algorithms

Algorithms for computing optical flow vary significantly in computational complexity, run-time and resulting accuracy. OpenCV library features several implementations of the optical flow estimation algorithms (as of version 3.4.1) and the algorithms tested here were: Lucas-Kanade sparse feature tracking [22], Gunnar Farneback’s algorithm of dense optical flow [23], the optical flow algorithm by Brox et. al. [24], Dual TV-Lı [25], Dense Inverse Search (DIS) [26], SimpleFlow [27], DeepFlow [28] and PCAFlow [29] (see Table 3).

**Table 3:** Tracking performance on the the simulated rotation dataset, using full frame optical flow.

| Algorithm | Mean magnitude<br>error, % | Mean orientation<br>error, degrees | Time,<br>ms |
| --- | --- | --- | --- |
| Dual TV-L1 (CUDA) | 4.9 | 1.95 | 953.2 |
| DeepFlow | 1.7 | 0.76 | 162.4 |
| Brox (CUDA) | 3.9 | 1.69 | 70.4 |
| PCAFlow (10x10 priors) | 1.6 | 0.73 | 33.1 |
| PCAFlow (default priors) | 2.0 | 0.84 | 26.4 |
| Farneback | 1.2 | 0.54 | 14.6 |
| Lucas-Kanade (CUDA) | 1.9 | 0.83 | 13.2 |
| Farneback (CUDA) | 1.2 | 0.54 | 10.5 |
| SparseToDense | 2.0 | 0.83 | 8.2 |
| DIS (medium) | 3.6 | 1.6 | 4.6 |
| DIS (fast) | 14.5 | 6.48 | 1.9 |
| DIS (ultrafast) | 18.7 | 8.29 | 0.5 |

The speedup through reducing the size of the ROI varies depending on the algorithm, which motivated a comparison of the run-times depending on the input size (Fig 12). To achieve a tracking frequency of 500 frames per second, the run-time should not exceed 2 ms, preferably be less to accommodate the frame preprocessing and ball motion estimation calculations. The algorithms relying on sparse features pre-selection fail in smaller ROIs (Sparse-to-dense, PCAFlow). Farneback, although being slow with default parameters, could be sped up significantly by reducing the number of scale levels and by limiting the internal iterations. It turned out to be the algorithm most suitable for the application, since it can handle a ROI of up to 60 × 140 px in under 2 ms when tuned for speed.

**Figure 11:**
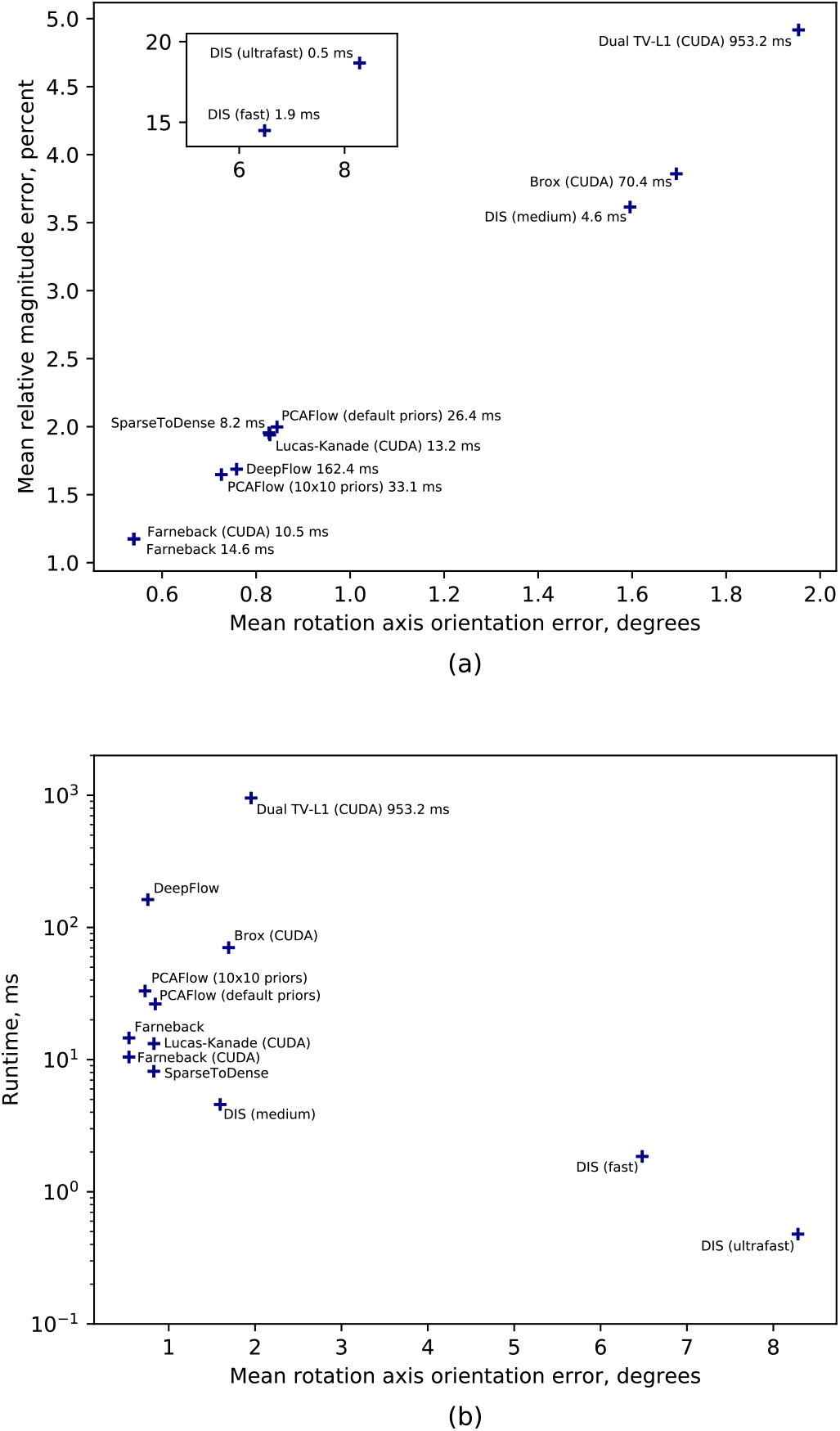
Ball rotation estimation errors (a) and run-time of the optical flow methods on full frame of the simulated dataset

**Figure 12:**
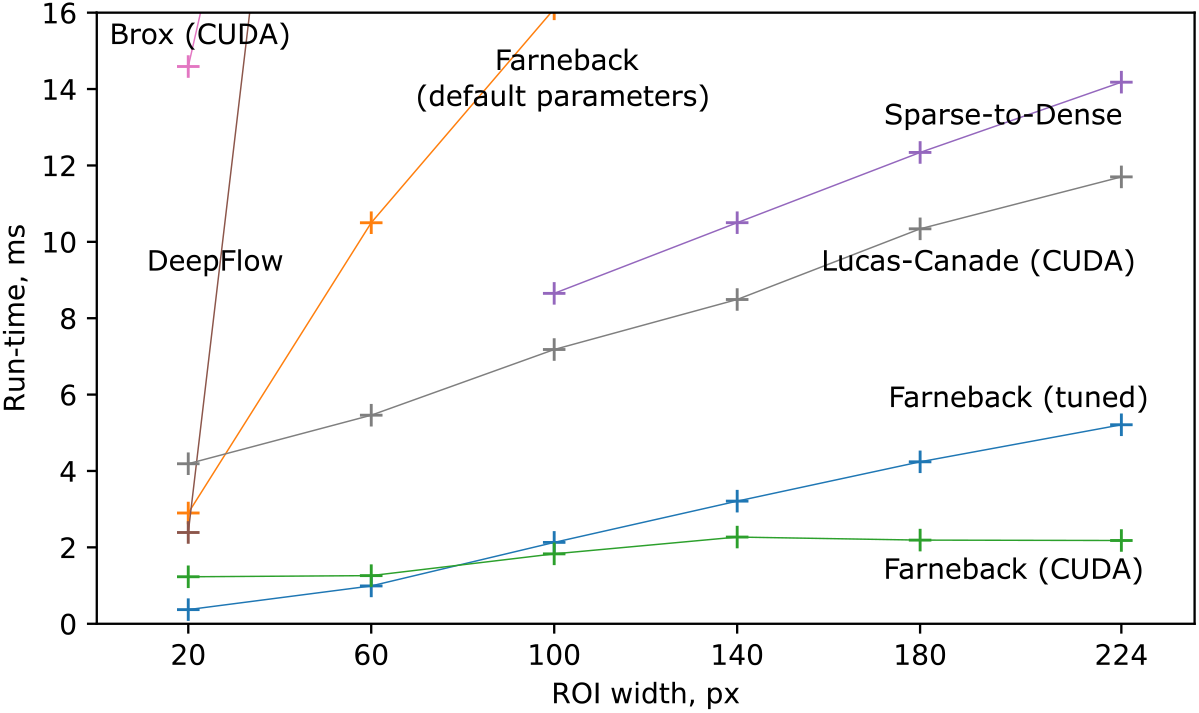
Run-time of optical flow algorithms depending on input size (ROI width × 140 px). Measured as average over 3000 frames with simulated dataset with accelerating rotations.

### 7.7. Selecting ROI parameters

The ROI is a small part of the full frame used to identify the rotation direction of the ball from optical flow. The ROI is limited by two concentric rings with their centers coinciding with the center of the frame, which is also selected to agree with the center of the tracked ball. By specifying the radius of the ROI *ρ_ROI_* (px) and its width Δ*ρ_ROI_* (px), one can vary the run-times of the fitting algorithm and its accuracy. The ROI parameters are chosen by running the tracking algorithm on simulated datasets (constant 1^°^ per frame angular velocity) and finding the variance in the estimated angular velocity magnitude and orientation compared to ground truth. Figure 13 shows the summed magnitude and orientation error (normalized) depending on *ρ_ROI_*, using Δ*ρ_ROI_* = 10 px. The lowest value was achieved for *ρ_ROI_* = 0.27, *ρ_max_* = 60 px.

**Figure 13:**
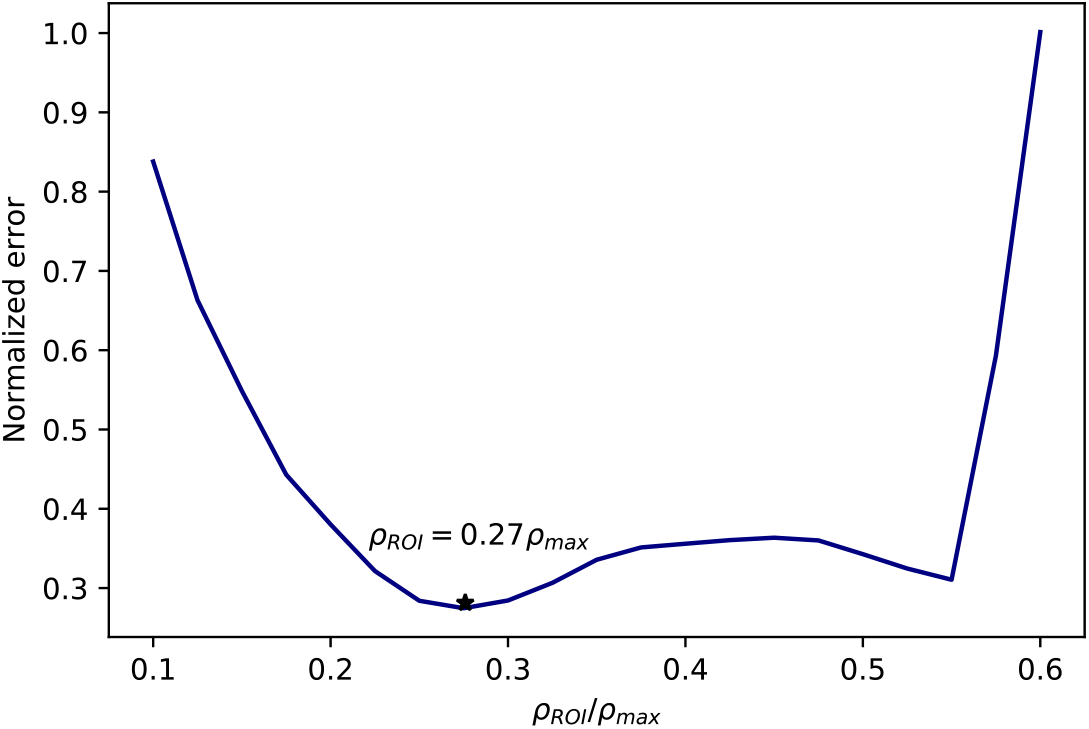
Summed tracking orientation and rotation errors (normalized), obtained by running the tracking algorithm with Δ*ρ_ROI_* = 10 px. *ρ_max_* is the visual radius of the ball (116 px).

### 7.8. Calibration coefficients

The coefficients *c_xy rad_, c_xy tan_* and *c_z_* in expression (8) are required for relating the optical flow in pixels (as determined with the optical flow algorithm) to the actual ball displacement in radians. Apart from a unit conversion factor these coefficients depend on the camera magnification, the size of the ball, and the position of the ROI on the ball. These factors can be determined in any of the following ways: by correlating optical flow induced by known ball rotations (with a ground truth dataset); similar, but using modelled optical flow with the pinhole camera model as in expressions (5-7, provided the optical parameters of the camera are known); or using a second synchronized camera. These methods are described in more detail in the following subsections.

#### 7.8.1. Calibration with ground truth dataset

The optical flow in the tracking ROI was calculated on simulated data and its distributions are fitted with function (8). Then the calibration constants can be found as the ratios:

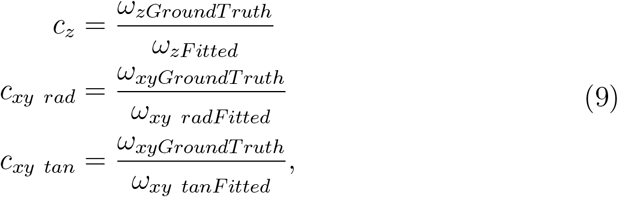

where *ω_xyGroundTruth_ ω_zGroundTruth_* are the known rotations, and *ω_xy radFitted_, ω_xy tanFitted_* are the results of fitting radial and tangential optical flow distributions in the ROI assuming *c_xy rad_* = *c_xy tan_* = *c_z_* = 1.0.

Using the rendered ball rotation dataset with constant rotations, the following calibration factors were found:

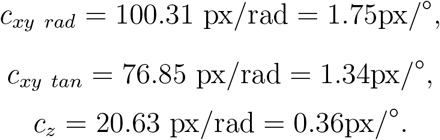

#### 7.8.2. Calculation of the calibration factors from the scene model

Using the ball projection model (equations 3 and 4) with substituted camera and scene parameters, optical flow distribution in the tracking ROI can be predicted and used for calibration similar to the previous method. With the help of a scaled-up calibration setup, the camera and scene parameters were measured as listed in Table 4.

**Table 4:**
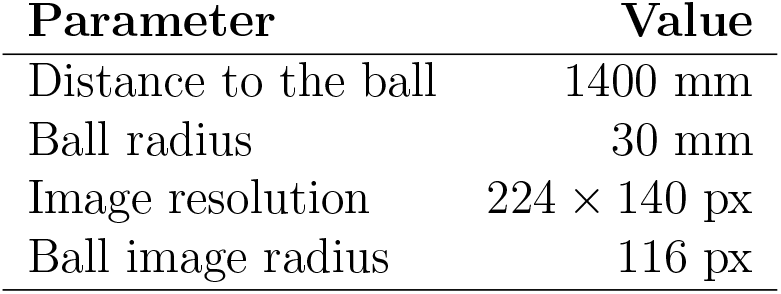
Measured parameters of the tracking setup scale model

Using these measurements, and the pinhole camera model (4) the focal length of the camera can be calculated (assuming the image projection center is exactly in the middle of the frame, *center_x_* =112 px, *center_y_* = 70 px):

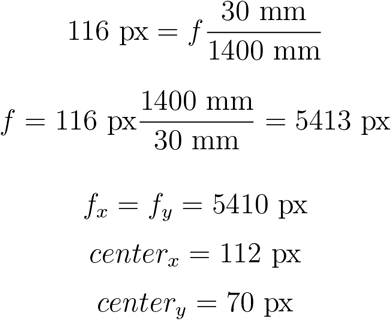

The model with these parameters was used to predict the optical flow induced by different rotations according to expressions ((5) – (7)).

The calibration factors found with this method were close to ones obtained with ground truth rotations:

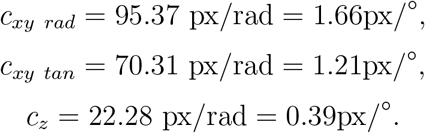

#### 7.8.3. Two-camera calibration

According to the mathematical model described above the z-component of the angular velocity measured by the tracking camera is detected as the xy-plane component by a second camera filming the same rotation from an orthogonal direction (see Figure 14).

**Figure 14:**
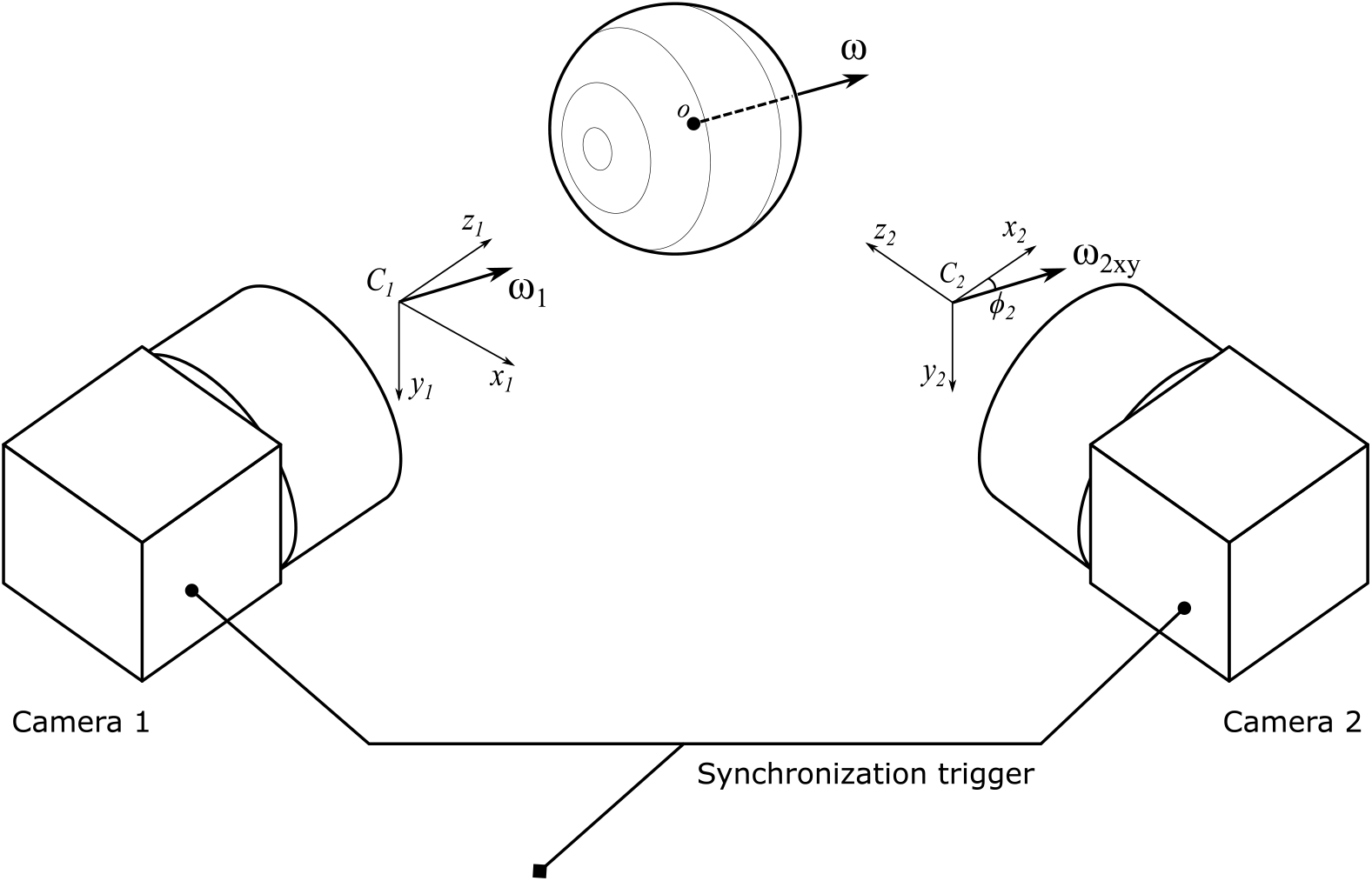
Calibration procedure using two cameras. The z-component of angular velocity of the ball in the reference frame of camera 1 is projected onto the x-axis in the reference frame of camera 2. Calibration is performed by correlating those components to find *c_xy_*.

As shown using the optical flow model (Figure 4), a rotation of the ball around the camera’s z-axis results in all of its points moving along circular trajectories in the frame plane. This means that the angular displacement of the ball equals the angular displacement of the points in the frame along their trajectories, and the tangential optical flow in the ROI induced by this motion is proportional to the radius of the ROI, meaning that *c_z_* can be estimated with a single camera without measuring any additionally parameters.

Different from z-factors, *c_xy rad_* and *c_xy tan_* depend on the camera properties and the scene geometry. Therefore, a second camera is introduced, filming the ball from an orthogonal direction, as shown in Figure (14). The z-component measured by camera 1 is then equal to the x-component measured by camera 2, and camera 1 can therefore be used to calibrate xy-tracking by camera 2:

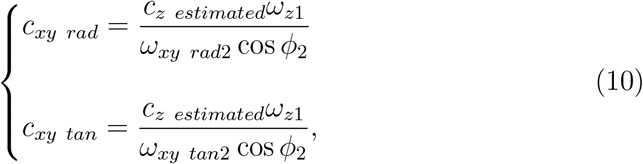

where *c_z estimated_* is estimated as described, *ω*_*z*1_ is measured by camera 1, *ω*_*xy rad*2_, *ω*_*xy tan*2_ and *ϕ*_2_ are found by fitting function (8) on radial and tangential optical flow in the ROI from camera 2, respectively, initially assuming *c_xy rad_* = *c_xy tan_* = 1.0. *ϕ*_2_ is the orientation of the ball’s axis of rotation in the xy-plane of camera 2, and the cosine yields the x-axis projection of that rotation, corresponding to measured z-rotation by camera 1.

To perform this calibration, both cameras need to be set up with identical objectives and aligned at the same distance from the tracked ball at right angles (see Figure 14); the ball is let to spin freely (at a speed within the range where the tracking algorithm is valid) with a stream of air while the cameras capture frames synchronized by an external triggering signal.

**Figure 15:**
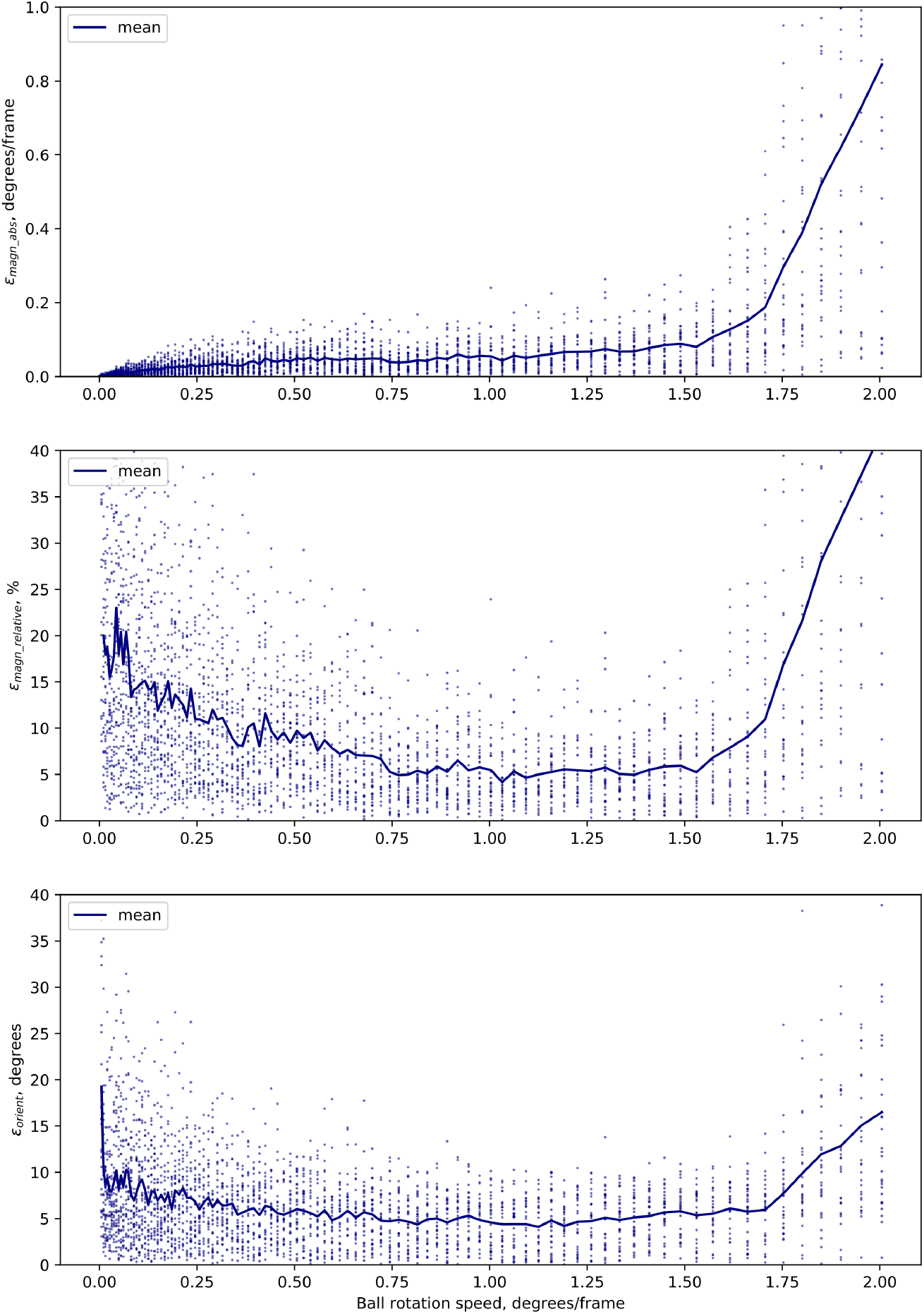
Tracking errors for an example of simulated, accelerated rotations, corresponding to the data shown in Figure 7.

## References

[1] D. A. Dombeck, M. B. Reiser, Real neuroscience in virtual worlds, Current opinion in neurobiology 22 (2012) 3–10.

[2] A. C. Mason, M. L. Oshinsky, R. R. Hoy, Hyperacute directional hearing in a microscale auditory system, Nature 410 (2001) 686.

[3] G. K. Lott, M. J. Rosen, R. R. Hoy, An inexpensive sub-millisecond system for walking measurements of small animals based on optical computer mouse technology, Journal of neuroscience methods 161 (2007) 55–61.

[4] D. A. Dombeck, A. N. Khabbaz, F. Collman, T. L. Adelman, D. W. Tank, Imaging large-scale neural activity with cellular resolution in awake, mobile mice, Neuron 56 (2007) 43–57.

[5] J. D. Seelig, M. E. Chiappe, G. K. Lott, A. Dutta, J. E. Osborne, M. B. Reiser, V. Jayaraman, Two-photon calcium imaging from head-fixed *Drosophila* during optomotor walking behavior, Nature methods 7 (2010) 535–540.

[6] R. J. Moore, G. J. Taylor, A. C. Paulk, T. Pearson, B. van Swinderen, M. V. Srinivasan, FicTrac: A visual method for tracking spherical motion and generating fictive animal paths, Journal of Neuroscience Methods 225 (2014) 106–119.

[7] H. Haberkern, M. A. Basnak, B. Ahanonu, D. Schauder, J. D. Cohen, M. Bolstad, C. Bruns, V. Jayaraman, On the adaptive behavior of head-fixed flies navigating in two-dimensional, visual virtual reality (2018).

[8] D. Turner-Evans, S. Wegener, H. Rouault, R. Franconville, T. Wolff, J. D. Seelig, S. Druckmann, V. Jayaraman, Angular velocity integration in a fly heading circuit, eLife 6 (2017) e23496.

[9] B. E. Bayer, An optimum method for two-level rendition of continuous tone pictures, in: IEEE International Conference on Communications, June, 1973, volume 26, 1973.

[10] Urho3D contributors, Urho3d: A cross-platform 2d and 3d game engine, https://urho3d.github.io/,2019.

[11] B. Palais, R. Palais, Eulers fixed point theorem: The axis of a rotation, Journal of Fixed Point Theory and Applications 2 (2007) 215–220.

[12] E. W. Weisstein, Spherical coordinates. From MathWorld-A Wolfram Web Resource, 2005. URL: http://mathworld.wolfram.com/SphericalCoordinates.html, visited on 20.03.2018.

[13] P. Sturm, Pinhole camera model, in: Computer Vision, Springer, 2014, pp. 610–613.

[14] G. Bradski, The OpenCV Library, Dr. Dobb’s Journal of Software Tools (2000).

[15] G. Bradski, Camera calibration and 3d reconstruction. From OpenCV Reference Manual, 2016. URL: https://docs.opencv.org/3.2.0/d9/d0c/group__calib3d.html, visited on 15.03.2019.

[16] J. A. Nelder, R. Mead, A simplex method for function minimization, The computer journal 7 (1965) 308–313.

[17] M. J. Grabowska, J. Steeves, J. Alpay, M. Van De Poll, D. Ertekin, B. van Swinderen, Innate visual preferences and behavioral flexibility in drosophila, Journal of Experimental Biology 221 (2018) jeb185918.

[18] J. R. Stowers, M. Hofbauer, R. Bastien, J. Griessner, P. Higgins, S. Farooqui, R. M. Fischer, K. Nowikovsky, W. Haubensak, I. D. Couzin, et al., Virtual reality for freely moving animals, Nature methods 14 (2017) 995.

[19] A. Jönsson, contributors, Angelscript, 2018. URL: https://www.angelcode.com/angelscript/, visited on 14.03.2019.

[20] Urho3D contributors, Urho3d scene model, 2019. URL: https://urho3d.github.io/documentation/1.7/_scene_model.html, visited on 14.03.2019.

[21] S. K. J. Simen Svale Skogsrud, Grbl, https://github.com/grbl/grbl, 2009.

[22] J.-Y. Bouguet, Pyramidal implementation of the affine lucas kanade feature tracker description of the algorithm, Intel Corporation 5 (2001) 4.

[23] G. Farnebäck, Two-frame motion estimation based on polynomial expansion, in: Scandinavian conference on Image analysis, Springer, 2003, pp. 363–370.

[24] T. Brox, A. Bruhn, N. Papenberg, J. Weickert, High accuracy optical flow estimation based on a theory for warping, in: European conference on computer vision, Springer, 2004, pp. 25–36.

[25] C. Zach, T. Pock, H. Bischof, A duality based approach for realtime tv-l 1 optical flow, in: Joint Pattern Recognition Symposium, Springer, 2007, pp. 214–223.

[26] T. Kroeger, R. Timofte, D. Dai, L. Van Gool, Fast optical flow using dense inverse search, in: European Conference on Computer Vision, Springer, 2016, pp. 471–488.

[27] M. Tao, J. Bai, P. Kohli, S. Paris, Simpleflow: A non-iterative, sublinear optical flow algorithm, in: Computer Graphics Forum, volume 31, Wiley Online Library, 2012, pp. 345–353.

[28] P. Weinzaepfel, J. Revaud, Z. Harchaoui, C. Schmid, Deepflow: Large displacement optical flow with deep matching, in: Computer Vision (ICCV), 2013 IEEE International Conference on, IEEE, 2013, pp. 1385–1392.

[29] J. Wulff, M. J. Black, Efficient sparse-to-dense optical flow estimation using a learned basis and layers, in: Proceedings of the IEEE Conference on Computer Vision and Pattern Recognition, 2015, pp. 120–130.

